# The Kelch 3 motif on gigaxonin mediates the interaction with NUDCD3 and regulates vimentin filament morphology

**DOI:** 10.1101/2025.03.10.641328

**Authors:** Cassandra L. Phillips, Christina So, Meredith F. Gillis, Jonathan Harrison, Chih-Hsuan Hsu, Diane Armao, Natasha T. Snider

## Abstract

Gigaxonin is an intermediate filament (IF)-interacting partner belonging to the Kelch-like (KLHL) protein family. Gigaxonin is encoded by the *KLHL16* gene, which is mutated in Giant Axonal Neuropathy (GAN). The lack of functional gigaxonin in GAN patient cells impairs IF proteostasis, leading to focal abnormal accumulations of IFs and compromised neuronal function. We hypothesized that gigaxonin forms molecular interactions via specific sequence motifs to regulate IF proteostasis. The goal of this study was to examine how distinct Kelch motifs on gigaxonin regulate IF protein degradation and filament morphology. We analyzed vimentin IFs in HEK293 cells overexpressing wild type (WT) gigaxonin, or gigaxonin lacking each of the six individual Kelch motifs: K1 (aa274-326), K2 (aa327-374), K3 (aa376-421), K4 (aa422-468), K5 (aa470-522), and K6 (aa528-574). All six gigaxonin deletion mutants (ΔK1-ΔK6) promoted the degradation of soluble vimentin. The ΔK3 gigaxonin mutant exhibited soluble vimentin degradation and promoted the bundling of vimentin IFs relative to WT gigaxonin. Using mass spectrometry proteomic analysis we found that, relative to WT gigaxonin, ΔK3 gigaxonin had increased associations with ubiquitination-associated and mitochondrial proteins and lost the association with the NudC domain-containing protein 3 (NUDCD3), a molecular chaperone enriched in the nervous system. Collectively, our cell biological data show the induction of an abnormal GAN-like IF phenotype in cells expressing ΔK3-gigaxonin, while our mass spectrometry profiling links the loss of gigaxonin-NUDCD3 interactions with defective IF proteostasis, revealing NUDCD3 as a potential new target in GAN.

## Introduction

Gigaxonin is an intermediate filament (IF)-interacting partner that belongs to the 42-member family of Kelch-like (KLHL) proteins (Bomont et al., 2000; Shi et al., 2019). KLHL proteins form diverse molecular interactions and regulate protein degradation and trafficking in different cell types (Dhanoa et al., 2013). Precise functioning of KLHLs is essential in the muscle and brain, which is evident by the physiological effects of disease-causing mutations within this gene family. *KLHL16* (or *GAN*) is the gene that encodes gigaxonin, and it is mutated in the pediatric neurodegenerative disease Giant Axonal Neuropathy (GAN) (Bharucha-Goebel et al., 2021). The lack of functional gigaxonin in GAN patient cells impairs IF proteostasis, leading to defective IF dynamics, focal abnormal accumulation of IF proteins, and compromised cellular function, especially in neurons (Johnson-Kerner et al., 2014). Molecular-level understanding of gigaxonin-IF interactions can provide new insights into IF network regulation and GAN pathogenesis.

Gigaxonin interacts with IF proteins via its ∼300 amino acid-spanning C-terminal Kelch domain (Johnson-Kerner et al., 2015). The Kelch domain is composed of six individual Kelch motifs (K1-K6), but the significance of this domain and its individual motifs on IF proteostasis is not clear. We observed that clinical variants of unknown significance (VUS) impacting the Kelch domain of gigaxonin have high predicted pathogenicity based on protein structure models (Cheng et al., 2023). With the aim of advancing knowledge of the functional and disease relevance of gigaxonin, we sought to determine how the deletion of each individual Kelch motif on gigaxonin affects its function.

To accomplish this, we compared soluble vimentin protein levels and vimentin IF organization in HEK293 cells overexpressing wild type gigaxonin or gigaxonin lacking each of the six Kelch motifs. Unexpectedly, our data revealed a disconnect between the levels of soluble vimentin (substrate for gigaxonin-induced degradation) and effects on filament morphology. We conducted mass spectrometry proteomics, which revealed new gigaxonin interactors, and uncovered a novel interaction between gigaxonin and the NUDCD3 chaperone. The findings point to a potential new mechanism behind the pathologic effect of gigaxonin deficiency and may help to explain high pathogenicity predictions from the structure-based analysis of gigaxonin variants affecting specific segments of the Kelch domain.

## Results

### Distribution of known and predicted pathogenic variants across the gigaxonin protein domains

Based on the genetic analysis from a GAN natural history study (Bharucha-Goebel et al., 2021), the majority of the pathogenic variants are gigaxonin missense mutations (**Fig. 1A**). The missense mutations are spread across the three gigaxonin domains (BTB, BACK, and Kelch) without any evidence for ‘hotspots’, and there is currently no known genotype-phenotype correlation (Bharucha-Goebel et al., 2021). The recently developed deep learning model AlphaMissense uses sequence-based structural protein models to assign pathogenicity scores for variants (Cheng et al., 2023). For missense variants reported in the natural history study of GAN patients (Bharucha-Goebel et al., 2021), this tool accurately predicted the high pathogenicity of 19/22 variants with median score of 0.9850, on a scale from 0 to 1 (**Fig. 1B**). Therefore, we applied AlphaMissense to all GAN coding missense variants classified as uncertain significance, likely pathogenic, or pathogenic in the NIH ClinVar database (Landrum et al., 2018) (**Fig. 1C**). Many GAN variants of unknown significance had high pathogenicity scores (red dots) that were similar to those of the known disease-causing variants (black dots). Frequency distribution analysis according to protein domain shows a higher number of predicted pathogenic variants map to the Kelch domain, especially variants with scores between 0.9 and 1.0 (**Fig. 1D**). The Kelch domain interacts with IF protein substrates, but how each individual segment of this domain contributes to overall gigaxonin function is not known. Therefore, we were interested in investigating the potential roles for the individual Kelch motifs.

**Figure 1.**
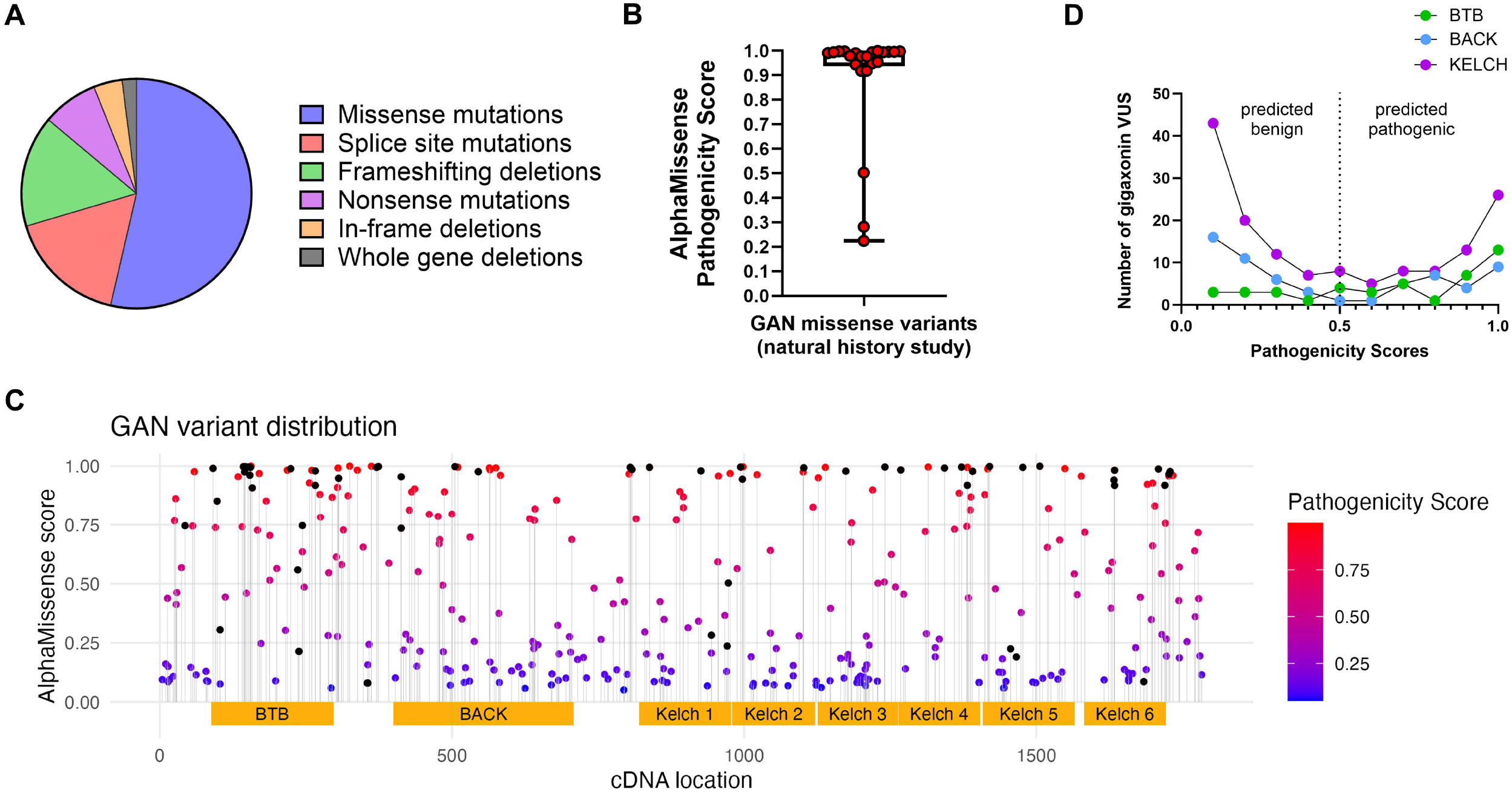
Pathogenicity of GAN missense variants across the gigaxonin protein domains. **A**. Missense mutations are the most common GAN variants; based on a genetic analysis of the GAN natural history study cohort in Bharucha-Goebel et al., *Brain* 2021. **B**. AlphaMissense scores on known disease-causing GAN variants from the natural history study, as represented by box-and-whisker plot (n=22; median score=0.9850). **C**. Missense GAN variants from ClinVar plotted by their respective cDNA location (x-axis) and AlphaMissense pathogenicity score (y-axis). Black dots represent known disease-causing GAN variants; from Bharucha-Goebel et al., *Brain* 2021 and Lescouzeres & Bomont, *Front Physiol* 2020. Blue and red dots represent variants of unknown significance (VUS), where blue is predicted benign and red is predicted pathogenic. Yellow boxes represent the gigaxonin functional domains BTB, BACK, and Kelch, as defined in UniProt (ID: Q9H2C0). **D**. Frequency distribution plot of AlphaMissense pathogenicity scores of gigaxonin VUS, according to the protein domain affected: BTB (green), BACK (blue), KELCH (magenta).

### Validation of a GFP-tagged human gigaxonin protein construct for molecular analysis of gigaxonin

We utilized a full-length human gigaxonin construct incorporating a C-terminal turbo-GFP (tGFP) tag (**Fig. 2A**). Both GFP-tagged (GFP-Gig) and untagged gigaxonin reduced soluble vimentin levels when transfected in HEK293 cells, showing that the GFP tag did not interfere with the vimentin degradation-promoting function of gigaxonin (**Fig. 2B-C**). To further validate that GFP-tagged gigaxonin can promote the degradation of other IF gigaxonin substrates, we examined glial fibrillary acidic protein (GFAP) in U251 cells transfected with GFP-gigaxonin. We found that GFP-Gig was active in promoting the degradation of GFAP (**Fig. 2D-E**). Using the validated GFP-tagged gigaxonin, we proceeded to analyze the impact of individual Kelch motif deletions on gigaxonin function.

**Figure 2.**
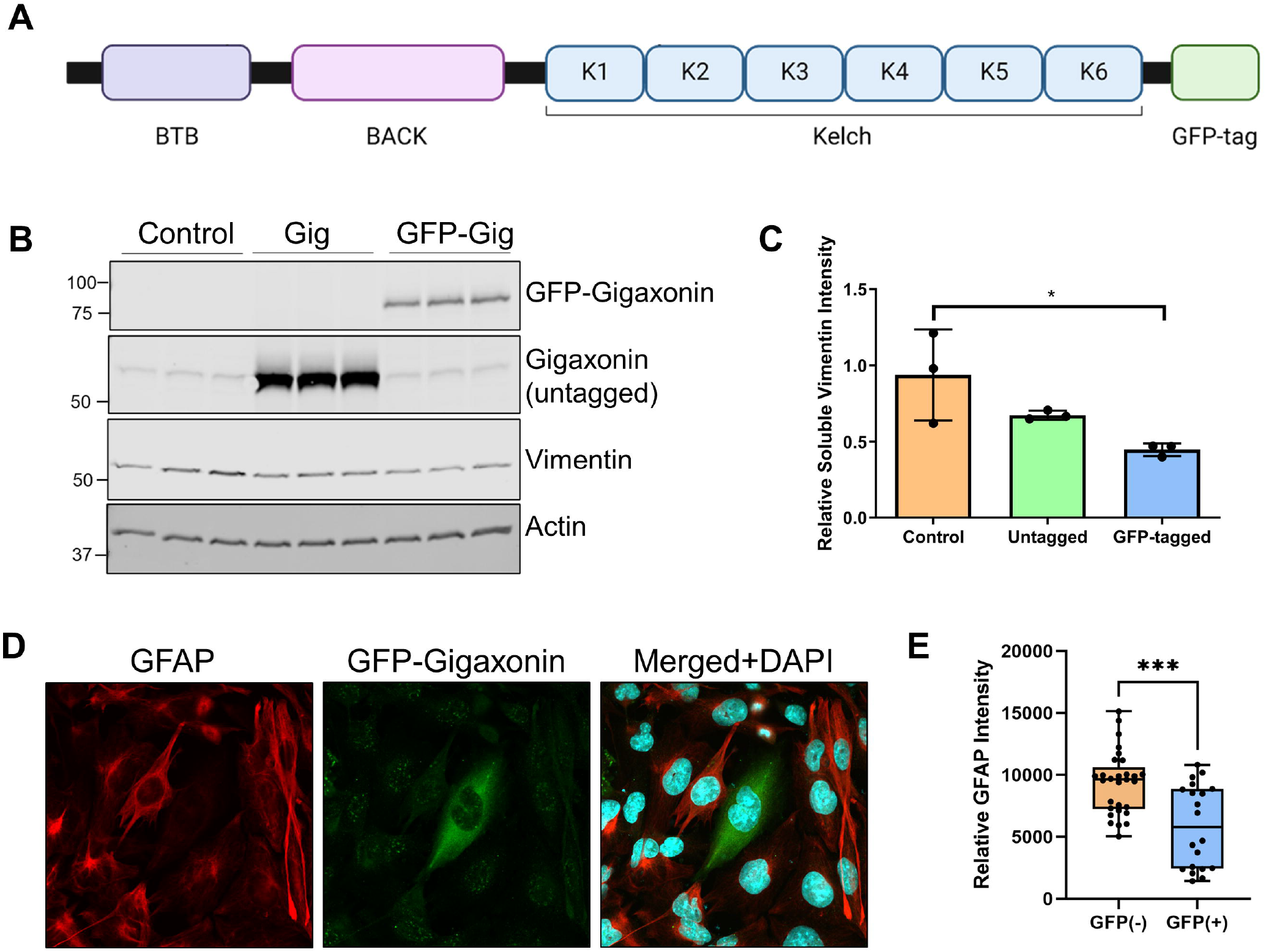
Functional validation of GFP-tagged wild type gigaxonin. **A**. Schematic representation of the gigaxonin domains and the location of the turbo-GFP tag at the C-terminus. **B**. Immunoblot for wild type GFP-tagged and untagged gigaxonin and the effect of their overexpression on TritonX-soluble vimentin levels in HEK293 cells; actin serves as a loading control. **C**. Quantification of the vimentin immunoblots from panel B. *p<0.05; one-way ANOVA. **D**. Overexpression of wild type GFP-gigaxonin (green) in U251 glioma cells and the effect on GFAP (red) clearance. ***p<0.001; unpaired t-test.

### Individual gigaxonin Kelch deletion mutants retain the ability to promote vimentin degradation

The six Kelch motifs on gigaxonin assemble into a beta-propeller structure that facilitates binding interactions with IF proteins and potentially other gigaxonin substrates and regulators (Boizot et al., 2014) (**Fig. 3A**). We generated constructs of gigaxonin lacking each of the six Kelch motifs: ΔKelch 1 (ΔK1; aa274-326), ΔKelch 2 (ΔK2; aa327-374), ΔKelch 3 (ΔK3; aa376-421), ΔKelch 4 (ΔK4; aa422-468), ΔKelch 5 (ΔK5; aa470-522), and ΔKelch 6 (ΔK6; aa528-574). Immunoblot analysis showed the expression of gigaxonin consistent with the presence of each deletion (**Fig. 3B**) and each construct revealed broad cytoplasmic gigaxonin distribution (**Fig. 3C**). To determine whether the IF degradation function of gigaxonin would be compromised by the removal of each individual motif, we compared TritonX-soluble vimentin protein levels by immunoblot and quantitative ELISA. Soluble vimentin was reduced similarly in the deletion mutants compared to WT-Gig (**Fig. 3D)**. Notably, the ELISA assay suggested that ΔK3-Gig may be more active in degrading vimentin compared to WT-Gig (**Fig. 3E**). Overall, these results show that the individual Kelch motifs are dispensable for the ability of gigaxonin to target soluble vimentin for degradation.

**Figure 3.**
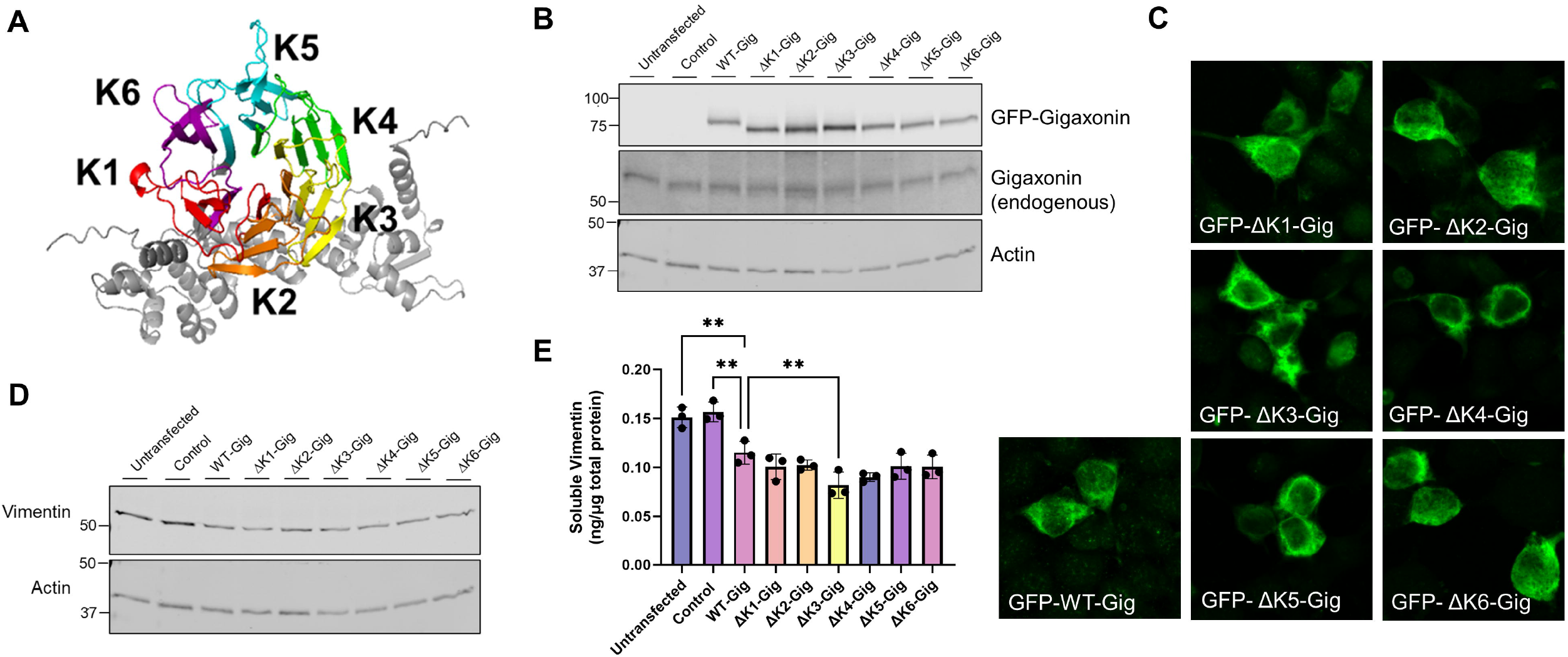
Effect of WT and Kelch deletion mutant gigaxonin expression on soluble vimentin. **A**. AlphaFold model of gigaxonin showing the predicted beta propeller structure and location of each Kelch motif that compose the gigaxonin Kelch domain. **B**. Immunoblot of GFP-gigaxonin and endogenous gigaxonin in cells expressing WT or Kelch deletion mutants. **C**. Immunofluorescence analysis of GFP-tagged WT and Kelch deletion mutants of gigaxonin. **D**. Immunoblot of TritonX-soluble vimentin in HEK293 cells transfected with WT and Kelch deletion mutants of gigaxonin. **E**. Quantitative vimentin ELISA of soluble vimentin. **p<0.01; one-way ANOVA compared to WT-Gig group.

### Deletion of the gigaxonin Kelch 3 motif promotes vimentin bundling

Next, we analyzed the morphology of vimentin IFs in HEK293 cells transfected with the WT gigaxonin and the Kelch deletion constructs. The organization of the vimentin filament network was similar across the transfections, with the exception of cells expressing ΔK3, where it appeared more bundled or aggregated (**Fig. 4**). Lipofectamine-only control cells contained thin and elongated vimentin filaments (**Fig. 5A**), while WT-Gig expressing cells had partial or full loss of the vimentin network (**Fig. 5B**). In contrast, ΔK3-Gig expressing cells displayed numerous thick vimentin bundles and aggregate-like structures (**Fig. 5C**). Particle analysis using ImageJ revealed a significant increase in the number and size of vimentin particles in ΔK3-Gig transfected cells compared to WT-Gig transfected cells (**Fig. 5D**). Since IF aggregation is a feature commonly observed in cells impacted by GAN-causing gigaxonin mutations (Mahammad et al., 2013), the data allude to the possibility that, despite being dispensable for degrading a pool of soluble vimentin, the Kelch 3 motif may be involved in other mechanisms that regulate the vimentin filament structure.

**Figure 4.**
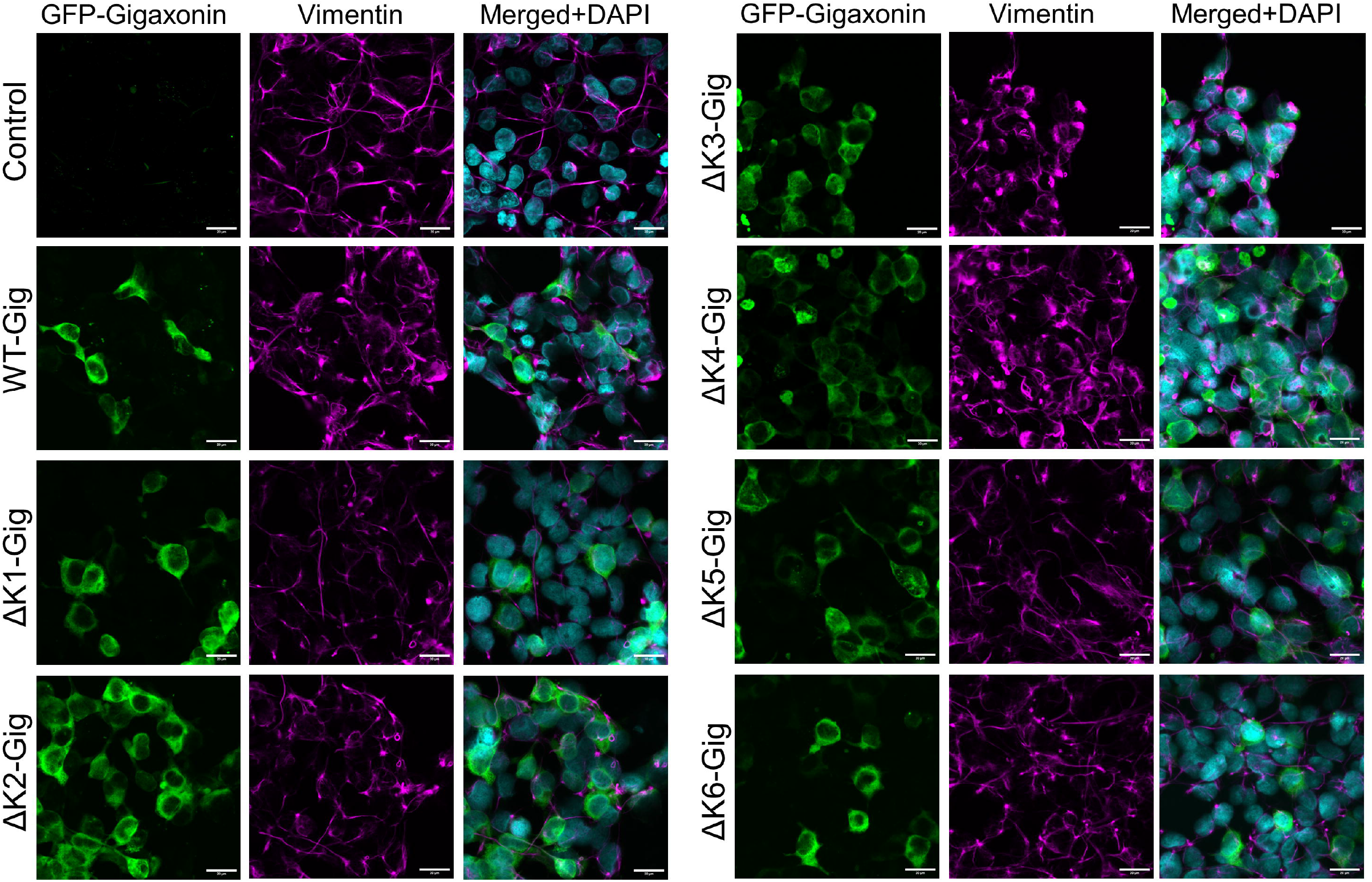
Vimentin filament morphology in HEK293 cells expressing WT and Kelch deletion mutants of gigaxonin. Immunofluorescence analysis of GFP-gigaxonin (green), vimentin (magenta), and DAPI (cyan) in HEK293 cells transfected with GFP-tagged WT gigaxonin (Gig) and Kelch motif deletion mutants (ΔK1-K6); Lipofectamine-only condition serves as transfection control. Scale bars=20μm.

**Figure 5.**
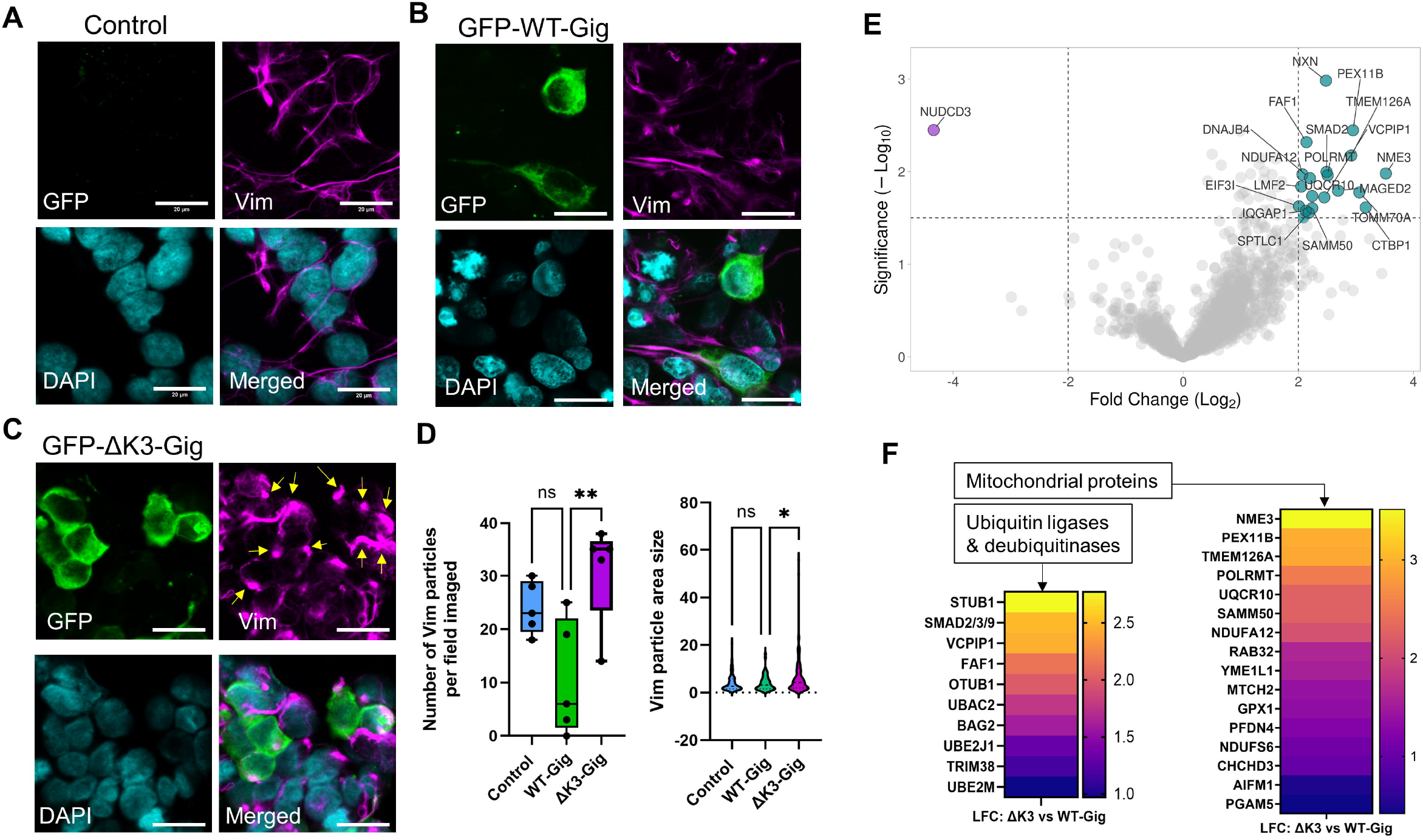
Effect of gigaxonin Kelch 3 deletion on vimentin filament morphology and gigaxonin protein interactions. **A-C**. Immunofluorescence analysis of GFP-gigaxonin (green), vimentin (magenta), and DAPI (cyan) in HEK293 cells treated with **(A)** Lipofectamine only or transfected with **(B)** WT or **(C)** Kelch 3 deletion mutant (ΔK3) GFP-gigaxonin. Yellow arrows point to vimentin bundles in ΔK3 condition. Scale bars=20μm. **D**. ImageJ analysis of number and area size of vimentin particles in Lipofectamine control (blue), WT-Gig (green), and ΔK3-Gig (magenta) conditions. **p<0.01; *p<0.05; one-way ANOVA compared to WT-Gig group. **E**. Volcano plot of protein interactors of ΔK3-gigaxonin that were significantly changed from WT-gigaxonin, based on mass spectrometry analysis on GFP-Gig pulldowns. Cyan dots represent increased interactions; magenta dot represents decreased interaction. **F**. Heat maps of the two major categories of differentially interacting gigaxonin partners in the absence of the Kelch 3 motif. Left map shows ubiquitin ligases/deubiquitinases and right map shows mitochondrial proteins (selected out of 71 statistically significant proteins detected in the ΔK3-Gig pulldown relative to WT-Gig).

### The Kelch 3 motif mediates gigaxonin protein interactions

Seeking to uncover new interactions of gigaxonin that are dependent upon the Kelch 3 motif, we conducted a mass spectrometry analysis on GFP pulldowns from cells overexpressing GFP-WT and GFP-ΔK3 gigaxonin. This analysis identified 71 interactions that were significantly changed by the K3 deletion (**Supplemental Table 1**). Deletion of K3 primarily led to increased association between gigaxonin and a number of chaperones, mitochondrial proteins, and ubiquitin-interacting proteins (**Fig. 5E-F**). Notably, increased interactions were observed with several ubiquitin ligases, including STUB1, UBE2M, UBE2J1, TRIM38 and the ubiquitin ligase regulator BAG2; deubiquitinases VCPIP1 and OTUB1; and the ubiquitin-binding proteins FAF1 and UBAC2 (**Fig. 5F**). A large number of mitochondrial proteins associated more strongly with ΔK3-Gig, most notably NME3, a known regulator of mitophagy and mitochondrial dynamics (**Fig. 5F**). Strikingly, the Kelch 3 deletion eliminated the association between gigaxonin and a single protein – the NudC domain-containing protein (NUDCD3) (**Fig. 5E**). NUDCD3 is a molecular chaperone that is enriched in the nervous system, associates with the cytoplasmic dynein complex, and regulates mitochondria motility. Using immunoblot analysis, we confirmed that the NUDCD3-gigaxonin interaction was eliminated by Kelch 3 deletion (**Fig. 6A**). AlphaFold prediction model revealed gigaxonin interacting with NUDCD3 via its Kelch domain (**Fig. 6B**), and this binding is eliminated by the removal of Kelch 3 (**Fig. 6C**). These data reveal a potential link between the loss of gigaxonin-NUDCD3 interactions with impaired IF proteostasis.

**Figure 6.**
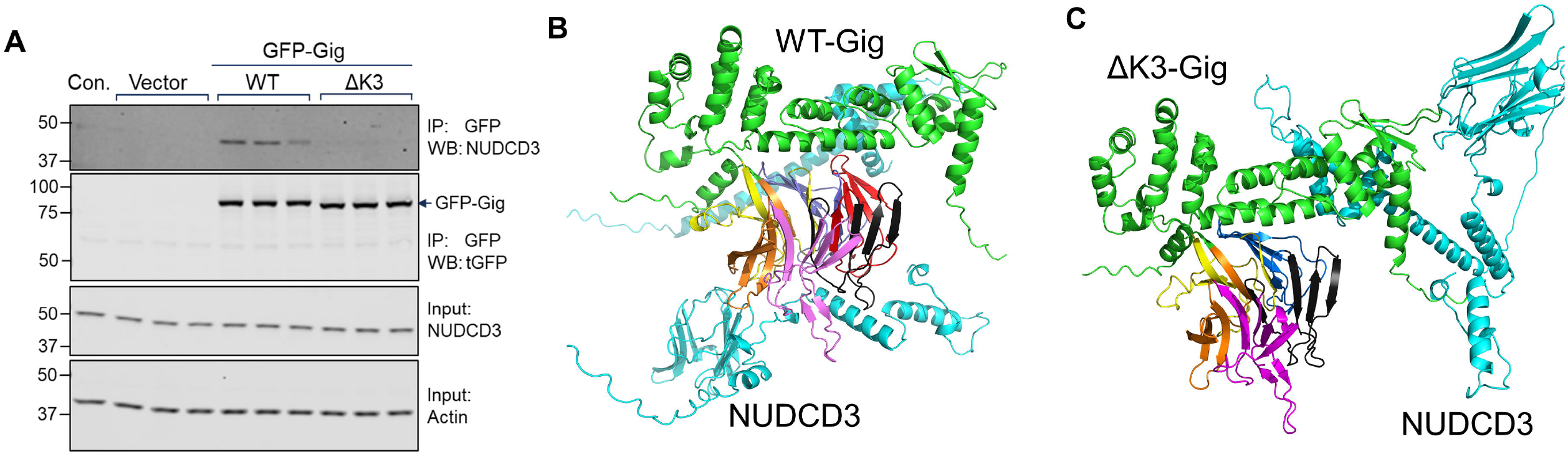
The Kelch 3 motif is required for gigaxonin interaction with the chaperone NUDCD3. **A**. Immunoblot analysis of NUDCD3 and GFP-Gig in GFP pulldowns on cell lysates from Lipofectamine control, GFP empty vector, and GFP-gigaxonin conditions (WT and ΔK3 mutant). Input lanes show total levels of NUDCD3 and actin (loading control) in the lysates. **B**. AlphaFold 3 predicted model of gigaxonin (green) interacting with NUDCD3 (cyan). Kelch repeat motifs are colored by domain: Kelch 1 (aa274-326, yellow), Kelch 2 (aa327-374, blue), Kelch 3 (aa376-421, red), Kelch 4 (aa422-468, black), Kelch 5 (470-522, purple), Kelch 6 (528-574, orange). **C**. AlphaFold 3 predicted model of gigaxonin ΔK3 (aa376-421) (green) interacting with NUDCD3 (cyan), with the remaining Kelch motifs colored as in **B**. With the removal of Kelch 3, NUDCD3 is no longer predicted to interact with the Kelch domain.

## Discussion

Our results revealed the importance of the gigaxonin Kelch 3 motif for the maintenance of proper protein interactions and vimentin filament structure. While it has been previously reported that the Kelch domain on gigaxonin interacts with IF proteins to promote proteasomal turnover (Johnson-Kerner et al., 2015), the contributions of the individual Kelch motifs to this degradation process have not been elucidated. Advancing knowledge of domain-specific functions for gigaxonin is essential to understand the high pathogenicity predictions for variants within the different protein segments, as this is translationally relevant to GAN disease-causing variants.

The functional significance of the interaction between gigaxonin and NUDCD3 requires further study. Our finding of gigaxonin/NUDCD3 interaction loss suggests a potential new disease mechanism associated with the gigaxonin deficiency present in GAN. Currently, there is limited mechanistic information about the molecular chaperone NUDCD3 (or NudCL). However, it has been reported to interact with both dynein-1 and dynein-2 motor proteins via the dynein intermediate chain (DIC) (Asante et al., 2014). In the absence of NUDCD3, HeLa cells exhibited impaired mitosis and subsequent cell death, dynein complex mislocalization, and DIC aggregation and degradation, suggesting an important role for NUDCD3 in maintaining normal dynein function in the context of cell proliferation (Zhou et al., 2006). Interestingly, overexpression of NUDCD3 was also shown to have detrimental effects on the cell cycle, resulting in compromised mitosis, cytokinesis, and cell division (Cai et al., 2009). Loss of NUDCD3 in retinal pigment epithelial (RPE) cells led to decreased expression of a DIC protein specific to the dynein-2 complex and reduced ciliogenesis, likely due to defective transport between dynein-2 and the cilia (Asante et al., 2014). Further, mutations in dynein-2-associated DIC proteins led to diminished interactions with NUDCD3, which in turn contributed to the destabilization of both the DIC and overall dynein complex (Vuolo et al., 2018). Inhibition of NUDCD3 partially reduced dynein-mediated retrograde transport of mitochondria in rat hippocampal neurons while knockdown of NUDCD3 in conjunction with another NUD family member led to a nearly complete loss of transport (Shao et al., 2013). It has been proposed that NUDCD3 functions as a chaperone to assist with proper folding of proteins with β-propellers, including numerous WD40-repeat proteins like the DIC proteins, and it has been reported to interact with other KLHL proteins (Taipale et al., 2014; Vuolo et al., 2018). Since the Kelch domain of gigaxonin is predicted to also form a β-propeller structure (Boizot et al., 2014), it is possible that NUDCD3 facilitates the proper folding and stabilization of the Kelch domain. Although we did not observe evidence of significant aggregation or degradation of the GFP-tagged ΔK3 gigaxonin construct, the loss of interaction with NUDCD3 may have led to variations in the folding pattern with downstream impacts on the roles for gigaxonin that are unrelated to IF protein turnover. Additional studies are needed to discern the binding mechanisms of NUDCD3 to gigaxonin in order to further perturb this association and study the resulting functional effects on gigaxonin. Interestingly, NUDCD3 is enriched in the nervous system (Fagerberg et al., 2014) and has been reported to interact with KLHL proteins that are highly expressed in the brain, involved in actin-binding and neuronal development (rather than participating in the UPS), and implicated in neurological conditions and brain tumors (Dhanoa et al., 2013; Shi et al., 2019). This highlights the importance of further studying this gigaxonin-NUDCD3 interaction in neurons, the most severely affected cell type in GAN, both under homeostatic and pathological conditions.

A recent study demonstrated that loss of gigaxonin led to impaired kinesin-1-mediated trafficking of vimentin in RPE cells and neurofilaments in dorsal root ganglion neurons, while transport of other cargo remained unaffected (Renganathan et al., 2023). Interestingly, direct binding of vimentin to kinesin-1 in gigaxonin knockout cells seemed to partly restore IF transport, suggesting that the interaction between IFs and kinesin is disrupted when there is a loss of gigaxonin (Renganathan et al., 2023). It is not currently known if kinesin-1 directly or indirectly binds to IFs, or if this interaction incorporates gigaxonin as well. While the dynein complex has also been previously shown to transport neurofilament IFs (Uchida et al., 2009), potential trafficking defects associated with this motor protein in the context of GAN currently remain unknown. Additional studies involving the effects of mutated forms of gigaxonin (missense or Kelch motif deletions), rather than a complete knockout, on kinesin or dynein mediated IF transport will help to elucidate whether defective IF trafficking is a result of gigaxonin’s role as an E3 ligase adaptor or another function. Since cells expressing ΔK3-Gig exhibited IF phenotypes reminiscent of those observed in GAN patient cells, it is possible that gigaxonin may play a larger role in IF transport mechanisms outside of regulating soluble IF protein turnover. However, the unexpected finding that deletion of the individual Kelch motifs did not impact soluble vimentin degradation suggests that mutated forms of gigaxonin may retain some functional activity when expressed in cells. In most GAN patients, gigaxonin is extremely low or absent, although the exact molecular mechanisms for gigaxonin loss in the case of missense mutations are not clear. Further studies are needed to discern if mutant forms of gigaxonin can be stabilized, for example by overexpressing NUDCD3 or another chaperone, and if they retain sufficient functional activity to degrade IFs in the most severely affected cell types.

Collectively, our data demonstrated a GAN-like IF phenotype in cells expressing Kelch 3 motif-deficient gigaxonin, as well as associated the loss of gigaxonin-NUDCD3 interactions with altered IF proteostasis, building upon recent findings that gigaxonin regulates IF transport (Renganathan et al., 2023). Further, these findings suggest a potential mechanism contributing to the pathologic effect of gigaxonin deficiency and may provide insights on the high pathogenicity scores associated with specific Kelch domain segments, as predicted by the structure-based analysis of gigaxonin variants.

## Materials and Methods

### Antibodies

The following primary antibodies and concentrations were utilized: mouse anti-tGFP (Origene, OT12H8, IF: 1:200-1:350), rabbit anti-Gigaxonin (Novus, NBP1-49924, WB: 1:250-1:500, IF: 1:100), rabbit anti-Vimentin (Cell Signaling Technology, D21H3, WB 1:1000, IF 1:200), rabbit anti-GFAP (DAKO; IF: Z0334), mouse anti-Actin (Santa Cruz, SPM161, WB 1:1000), and mouse anti-NUDCD3 (Santa Cruz, WB: 1:200). The following secondary antibodies and concentrations were utilized: IRDye 800CW goat anti-rabbit IgG (LI-COR, WB 1:5000), IRDye 680RD donkey anti-mouse IgG (LI-COR, WB 1:5000), and Alexa 488-, Alexa 568-, and Alexa 594-conjugated goat anti-mouse and anti-rabbit antibodies (Invitrogen, IF 1:500).

### Site-directed mutagenesis, transformation, and preparation of pCMV6-AC-GFP vector, wild type gigaxonin, and gigaxonin Kelch motif deletion constructs

Mutagenesis of GFP-tagged wild-type gigaxonin (pCMV6-AC-GFP vector; Origene) was conducted with the In-Fusion® Snap Assembly kit (Takara Bio) in accordance with manufacturer’s instructions to generate the following domain deletions (see table below for primers designed using primer design tool from Takara Bio): Kelch 1, Kelch 2, Kelch 3, Kelch 4, Kelch 5, and Kelch 6. Transformation for GFP-tagged gigaxonin wild type and Kelch 1-6 deletion constructs was conducted using Stellar Competent Cells (Takara Bio) provided by the In-Fusion® Snap Assembly kit. The cell suspension from the transformation was spread onto an ampicillin agar plate, then incubated overnight at 37°C. Colonies were individually picked and inoculated overnight at 37°C shaking at 225 RPM in LB media (Fisher Scientific) with ampicillin (100µg/µL). Plasmid DNA was extracted using the PureLinkTM Quick Plasmid Miniprep kit (Invitrogen) and submitted for Sanger sequencing to check for off-target changes in the vector, coding region of gigaxonin, or GFP-tag. After confirming the sequences, previously inoculated cell suspension was added to LB media with ampicillin (100µg/µL), then incubated overnight at 37°C shaking at 225 RPM. Plasmid DNA was extracted using either the ZymoPURE II Plasmid Midiprep Kit (Zymo) or the QIAfilter Plasmid Midi Kit (Qiagen) to generate 100ng/µL and 500ng/µL transfection stocks. Additional transformations for the pCMV6-AC-GFP vector (Origene), untagged wild type gigaxonin (pCMV6-XL4 vector; Origene), and GFP-tagged gigaxonin wild type (pCMV6-AC-GFP vector; Origene), Kelch 3 deletion, and Kelch 4 deletion constructs was conducted using XL1-blue supercompetent cells (Aligent Technologies), which were utilized in accordance with product instructions. Plasmid DNA was extracted to generate 100ng/µL and 500ng/µL transfection stocks as described.

### Site-directed mutagenesis primers

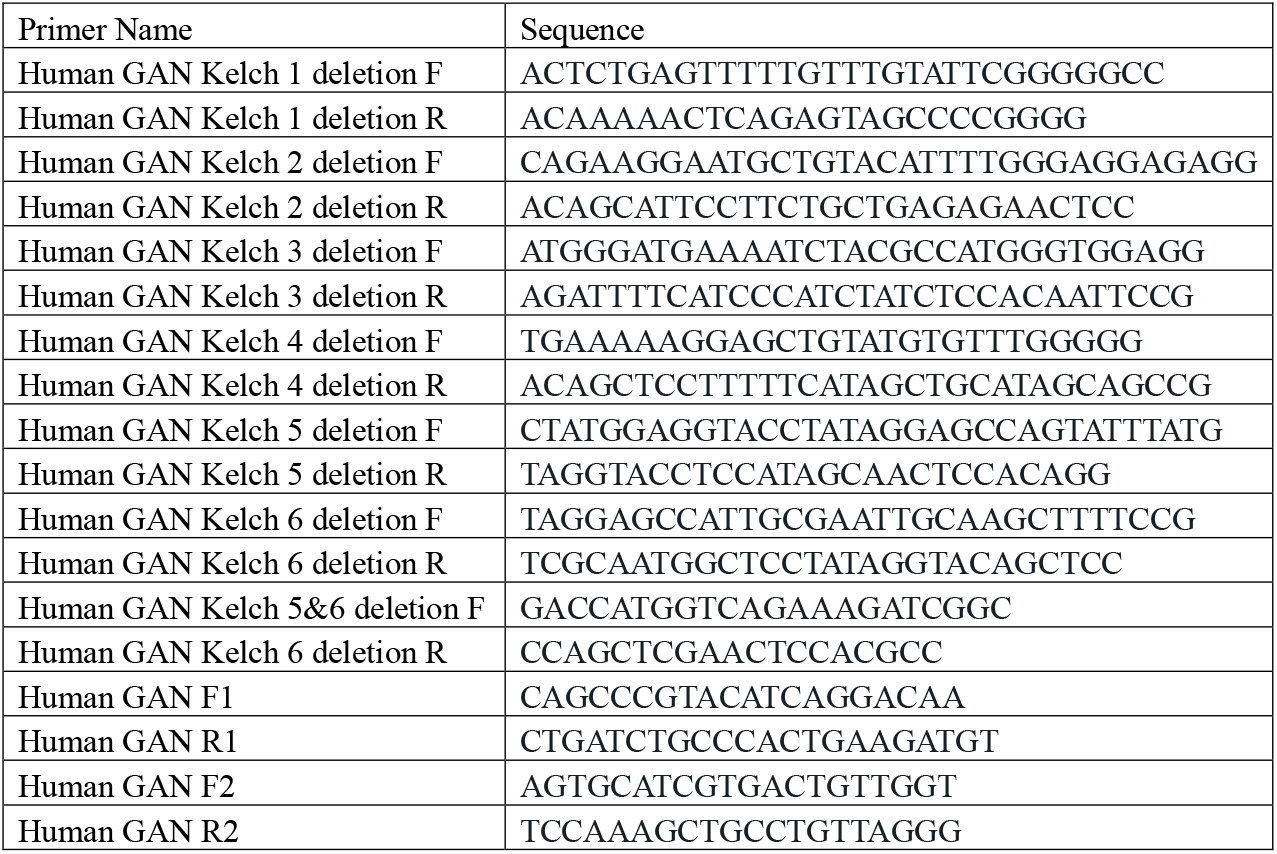

### Cell lines, cell culture, and transfection of gigaxonin mutants

HEK293 cells were cultured in MEM (ThermoFisher Scientific) with 10% FBS (Genesee Scientific, Lot: P093156 & Lot: P121400) and 1% penicillin/streptomycin (pen/strep; ThermoFisher Scientific). U251 cells were cultured in DMEM (ThermoFisher Scientific) with 10% FBS and 1% pen/strep. The media for both cell lines was changed every 2-3 days and cells were passaged at <95% confluency with 0.05% or 0.25% Trypsin-EDTA (ThermoFisher Scientific).

HEK293 cells or U251 cells were plated on 10cm plates or 8-well chamber slides at about 40-60% confluency in MEM +10% FBS media (no antibiotics). The following day, the cells were transfected with the pCMV6-AC-GFP vector, gigaxonin wild type construct, and/or the Kelch deletion constructs plus Lipofectamine 3000 and p3000 reagents that were utilized in accordance with product instructions (ThermoFisher Scientific). A Lipofectamine/p3000 only condition was utilized as well as an untransfected control. Media was changed to MEM +10% FBS + pen/strep media about 4-6 hours after transfection, and cells were harvested for immunoprecipitation (see below for details) or fixed for imaging (see below for details on immunostaining procedures) at the 48-hour time point.

For transfection experiments for immunoblot analysis, HEK293 cells were plated on 6-well plates at 80% confluency in MEM +10% FBS media (no antibiotics) with transfection reagents, including gigaxonin wild type and Kelch motif deletion plasmids, p3000, and Lipofectamine 3000. Media was changed to MEM +10% FBS + pen/strep media about 4-6 hours after transfection, and cells were harvested in TritonX-100 buffer at the 72-hour time point and separated into TCL, detergent soluble, and detergent insoluble fractions (see below for details on preparation of protein lysates).

### Preparation of protein lysates and immunoblotting

For immunoblot analysis, transfection lysates were prepared by adding 300-600µL cold TritonX-100 buffer (1% Triton X-100, 0.5 M EDTA, PBS, ddH_2_O) with protease/phosphatase inhibitors (Halt™ Protease & Phosphatase Inhibitor Cocktail, EDTA-free, 100X; ThermoFisher Scientific), scraping the cells with the plates on ice, and transferring the cell suspension into Eppendorf tubes; 10% of cell suspension was reserved as the TCL fraction and resuspended in an equal volume of hot 2X Novex™ Tris-Glycine SDS Sample Buffer (Thermo Fisher Scientific). The cell suspension was spun down in a tabletop centrifuge at 14,000xg for 10 min at 4°C to separate the detergent soluble and insoluble fractions, the supernatant was transferred to a new tube (detergent soluble fraction), then the remaining pellet (detergent insoluble fraction) was resuspended in hot sample buffer. All samples resuspended in with SDS sample buffer were heated at 95°C for 5 minutes and reduced with 5% 2-mercaptoethanol (Sigma) as needed. Total protein concentration for the detergent soluble fractions were measured with Pierce™ BCA Protein Assay Kit (Thermo Fisher Scientific) and Biotek Synergy HT plate reader according to manufacturer’s instructions; total protein concentrations were calculated from the recorded absorbance values. The concentrations were used to calculate 300ng/µL stocks; the samples were diluted with TritonX-100 buffer (+protease/phosphatase inhibitors). For immunoblotting, the 300ng/µL stocks were used to make 7.5ng stocks by adding an equal volume of hot SDS sample buffer to each sample, reducing with 5% 2-mercaptoethanol, and heating at 95°C for 5 minutes.

For immunoblotting, samples were loaded with equal volumes (resulting in 7.5ng per well) and separated on 10% or 4-20% Novex™ WedgeWell™ Tris-Glycine gels (Thermo Fisher Scientific) for 35-40 minutes at 225V or 1 hour 20-40 minutes at 120V and transferred at 40V overnight at 4°C onto nitrocellulose membranes. Gels were stained with Coomassie and destained following each transfer to verify normalization. The membranes were blocked in 5% non-fat milk (NFM) dissolved into 0.1% tween 20/PBS (PBST) at room temperature for 30-60 minutes. The membranes were incubated in primary antibodies diluted in 5% NFM/PBST at room temperature for 1 hour (see concentrations above), then washed 3x with PBST for 5 minutes each. The membranes were incubated with secondary antibodies diluted in 5% NFM/PBST at room temperature for 1 hour (see concentrations above), washed 3x with PBST and 1x with PBS for 5 minutes each, then scanned with a LI-COR Odyssey CLx machine.

### Immunoprecipitation with anti-GFP magnetic beads

For the immunoprecipitation (IP) experiment, transfection lysates were washed with HBSS and lysed in 1 mL cold TritonX-100 buffer (1% Triton X-100, 0.5 M EDTA, PBS, ddH_2_O) with protease/phosphatase inhibitors (Halt™ Protease & Phosphatase Inhibitor Cocktail, EDTA-free, 100X; ThermoFisher Scientific). The cell suspension was spun down in a tabletop centrifuge at 14,000xg for 20 min at 4°C to separate the detergent soluble and insoluble fractions, and the supernatant was transferred to a new tube (detergent soluble fraction). Total protein concentration for the detergent soluble fractions were measured with Pierce™ BCA Protein Assay Kit (Thermo Fisher Scientific) and Biotek Synergy HT plate reader according to manufacturer’s instructions; total protein concentrations were calculated from the recorded absorbance values. The concentrations were used to calculate 1µg/µL stocks; the samples were diluted with TritonX-100 buffer (+protease/phosphatase inhibitors). To obtain an input immunoblot for comparison to the IP, a fraction of each 1µg/µL stock was resuspended with an equal volume of hot SDS sample buffer, reducing with 5% 2-mercaptoethanol, and heating at 95°C for 5 minutes. Samples were loaded into gels at equal volumes and SDS-PAGE/immunoblot procedures were conducted as described above.

ChromoTek TurboGFP-Trap Magnetic Agarose beads (Proteintech) were utilized for immunoprecipitation (IP) of GFP-tagged vector, gigaxonin wild type, and gigaxonin Kelch 3 and 4 deletion constructs in accordance with product instructions. In brief, the beads were equilibrated with 3x washes with TritonX-100 buffer (1% Triton X-100, 0.5 M EDTA, PBS, ddH2O) with protease/phosphatase inhibitors (Halt™ Protease & Phosphatase Inhibitor Cocktail, EDTA-free, 100X; Thermo Fisher Scientific); washes were manually removed when beads were separated using a magnetic rack. Following equilibration, detergent soluble 1µg/µL stocks were loaded onto magnetic beads and incubated overnight at 4°C while rotating. Following incubation, the supernatant was removed, the beads were washed 3x with TritonX-100 buffer (no inhibitors), then 20% of the IP was transferred for elution with hot SDS sample buffer and heated at 95°C for 10 minutes. These samples were loaded into gels at equal volumes and SDS PAGE/immunoblot procedures were conducted as described above. The remaining IP was washed 3x with 50mM Ammonium Bicarbonate (pH 7.8), then the captured protein was eluted in 50mM Ammonium Bicarbonate (pH 7.8) for mass spectrometry-based proteomics analysis (described below).

### Immunofluorescence staining and imaging

HEK293 cells or U251 cells were rinsed with PBS and fixed in methanol at -20°C for 15 minutes, washed 2x with PBS for 5 minutes each, and incubated in blocking buffer (2.5% Bovine Serum Albumin (Sigma), 2% normal goat serum (ThermoFisher Scientific), PBS) at room temperature for 1 hour. Cells were incubated with primary antibodies at room temperature for 1-2 hours or overnight at 4°C, followed by 3x 5-minute PBS washes, then incubated with Alexa Fluor-conjugated secondary antibodies at room temperature for 1 hour and washed 3x with PBS for 5 minutes each. Finally, cells were incubated in DAPI (Invitrogen), washed 2x with PBS for 5 minutes each, and mounted in Fluoromount-G (SouthernBiotech) overnight. Cells were imaged on Zeiss 880 confocal laser scanning microscope using a 63x oil immersion objective.

### Quantitative vimentin ELISA

TritonX-100 detergent soluble fractions for HEK293 cells transfected for 72 hours with the wild type gigaxonin and Kelch deletion constructs, as well as untransfected and Lipofectamine3000 control samples, were analyzed using a vimentin ELISA (Cell Signaling Technology; Cat: 87105). The ELISA did not come with vimentin standards, so purified recombinant vimentin (gift from Dr. Harald Herrmann) of a known concentration was used and serially diluted to generate a standard range (in ng; 0.015, 0.03, 0.06, 0.12, 0.24, 0.49); standard curve concentrations were optimized to fall within the absorbance values measured by BioTek Cytation 5 plate reader. The 300ng/µL stocks were utilized to generate 50ng/µL samples (sample concentrations optimal for the standard curve were optimized in prior experiments), and the vimentin ELISA was performed in accordance with the manufacturer’s instructions. The absorbance values for each sample were averaged across three technical replicates, then vimentin concentration (in ng) was calculated based on a standard curve and normalized to total protein concentration per sample (in µg).

### Mass spectrometry-based proteomics analysis

GFP-tagged gigaxonin immunoprecipitation samples were prepared as described above and stored in 50mM Ammonium Bicarbonate (pH 7.8), then submitted for mass spectrometry-based proteomic analysis. In brief, on-bead tryptic digestion was performed, followed by peptide extraction and C18 desalting cleaning. Each sample was analyzed by LCMS/MS using the Thermo Easy nLC 1200-QExactive HF. The data were processed using MaxQuant (v1.6.15.0) – the data were searched against the Uniprot Human database (∼20,000 sequences), a common contaminants database (∼250 sequences), and gigaxonin sequences. The MaxQuant results were filtered and analyzed in Perseus, and reverse hits and proteins with 1 peptide (single hits) were removed. Perseus was used for imputation, log2 fold change and p-value calculations.

### Data analysis and statistics

AlphaMissense pathogenicity scores (Cheng et al., 2023) were obtained for all *KLHL16/GAN* gene coding missense variants classified as uncertain significance, likely pathogenic, or pathogenic in the NIH ClinVar database (Landrum et al., 2018) and for the variants reported from patient cohorts (Bharucha-Goebel et al., 2021; Lescouzeres and Bomont, 2020). Variants were plotted by cDNA location and AlphaMissense pathogenicity score with each dot representing a *KLHL16/GAN* gene variant based on cDNA change. The dots are color-coded by pathogenicity score with blue as predicted to be benign and red predicted as pathogenic, and the black dots are variants that were found in a patient cohort. Yellow boxes represent the functional domains of gigaxonin (BTB, BACK, and Kelch 1-6 motif repeats) as defined by UniProt (UniProt, 2023) (entry code for human gigaxonin: Q9H2C0). Additional graphical data were generated with GraphPad Prism.

Image Studio version 5.2 (LI-COR) was used to perform densitometry on the immunoblot for soluble vimentin, and the relative intensities were analyzed by the GraphPad Prism software using one-way ANOVA. The ELISA results for the untransfected, Lipofectamine3000, wild type gigaxonin, and Kelch 1-6 deletion conditions were analyzed using one-way ANOVA via GraphPad Prism. The mean relative GFAP intensities associated with untransfected cells and wild type GFP-tagged gigaxonin-transfected cells were measured using ImageJ and analyzed using one-way ANOVA via GraphPad Prism (note: additional experimental groups included in analysis are not included in this study). The number and size (micron^2^) of vimentin particles in the Lipofectamine3000, wild type gigaxonin, and ΔK3-gigaxonin conditions were measured using ImageJ via the analyze particles feature and analyzed using one-way ANOVA via GraphPad Prism. All graphical data were generated with GraphPad Prism. The proteomics data were analyzed using the DAVID (Sherman et al., 2022) bioinformatics resource, bar graphs and heat maps were generated with the GraphPad Prism software, and statistical analysis was performed via unpaired t-test.

## Summary of Supporting Information

- **Supplemental Table 1:** Significant interactors compared between wild type and Kelch 3 deletion GFP-tagged gigaxonin

## Acknowledgments

The authors thank Dr. Laura Herring and Dr. Allie Mills for assistance with the mass spectrometry analysis.

## Funding and Additional Information

This study was funded by Hannah’s Hope Fund and NIH grants R21NS121578 (NTS, DA), GM122741 (training award to CP), and AHA grant 24PRE1193707 (training award to CP). The content is solely the responsibility of the authors and does not necessarily represent the official views of the National Institutes of Health and the other sponsors.

## Author Contribution Statement

CLP: Conceptualization, Methodology, Validation, Formal analysis, Investigation, Visualization, Writing - Original Draft, Writing - Review & Editing, Funding acquisition; CS: Conceptualization, Methodology, Investigation, Writing - Review & Editing; MFG: Methodology, Formal analysis, Visualization, Writing - Review & Editing; JH: Methodology, Formal analysis, Visualization, Writing - Review & Editing; CHH: Investigation, Formal analysis, Visualization, Writing - Review & Editing; DA: Writing - Original Draft, Writing - Review & Editing, Supervision, Funding acquisition; NTS: Conceptualization, Methodology, Formal analysis, Visualization, Writing - Original Draft, Writing - Review & Editing, Resources, Supervision, Project administration, Funding acquisition

